# Physicochemical Feature-Enhanced Bipartite Graph Model for TCR-Epitope Binding Prediction with Tutorial Learning

**DOI:** 10.64898/2026.11.23.768277

**Authors:** Kenan Menguc, Alper Yilmaz

**Affiliations:** Industrial Engineering, Istanbul Technical University, Beşiktaş/Istanbul/ Turkey; Department of Civil, Environmental and Geodetic Engineering, The Ohio State University Columbus, Ohio 43210

**Keywords:** Graph Neural Networks (GNN), SMOTE, TCR–Epitope Binding, Bipartite Graphs, Tutorial Learning, Amino Acid Features

## Abstract

Accurate prediction of T-cell receptor (TCR) and epitope binding plays a key role in rapid immunological assessment, vaccine selection, and data-driven therapeutic design. However, existing computational approaches typically rely on primary amino acid sequences or NLP-based representations, which inadequately capture physicochemical interactions and structural relationships. Moreover, negative samples used for model training are frequently generated randomly, limiting discriminative learning and introducing considerable bias. This study proposes a feature-enriched bipartite graph architecture that incorporates multiple physicochemical properties to represent amino acids and sequence positions in a unified framework. Unlike methods that rely solely on primary sequence, our approach encodes residue-level physicochemical features that collectively influence three-dimensional conformation and binding interfaces. In addition, a tutorial-learning strategy is introduced to enrich model training with informative synthetic negatives without modifying model complexity. Experiments conducted on five epitope-specific datasets demonstrate up to a 5% improvement in predictive accuracy over state-of-the-art methods. We demonstrate epitope-specific model configurations as a practical approach for handling diverse binding scenarios, with a proposed pathway toward a generalizable multi-epitope framework. These results suggest that the proposed design offers a promising direction for computational immunology, particularly in low-data or highly imbalanced scenarios. Furthermore, an enhanced model design is conceptually introduced to accommodate future expansion of available epitope databases.

## 1 INTRODUCTION

T-cells are essential to the human immune system (Germain, 2002; Waldman et al., 2020; Wen et al., 2011). Antibodies neutralize and eliminate external viruses, protecting against infection (Cutillo et al., 2020). However, antibody production is initiated upon the detection of an abnormality (Graus et al., 2020). This process is regulated by the interaction between peptides presented on cell surfaces by major histocompatibility complex (MHC) molecules and T-cells. Specific antigen regions, termed epitopes, are recognized and targeted by the immune response (Pishesha et al., 2022). These epitopes are located in hydrophilic regions, thus increasing accessibility to the immune system.

Misrecognition of antigens by T-cell receptors (TCRs) can lead to autoimmune diseases, such as Type 1 diabetes, characterized by the attack on pancreatic beta cells, or cancer, attributable to neoantigen interactions and tumor-induced MHC suppression (Clark et al., 2021; Paucek et al., 2019). T-Cell Receptor - peptide Major Histocompatibility Complex (TCR-pMHC) binding, which entails intricate protein-protein interactions and co-receptor dynamics, is essential for immune regulation and response, with hypervariable regions like CDR3 playing a critical role in epitope recognition (Smirnov et al., 2023). A comprehensive understanding of these interactions is crucial for the development of personalized treatments and the advancement of drug discovery. Open-source databases, including VDJdb, McPAS, and IEDB, offer valuable resources for predictive model development in computational immunology, thereby supporting this vital area of research (Smirnov et al., 2023)

However, current computational methods have limitations, leading to our main motivation is to develop a feature-rich (physicochemical aware) learning framework for TCR–epitope binding prediction that moves beyond primary-sequence representations and conventional NLP-based approaches. Unlike methods that rely solely on primary sequence, our approach encodes residue-level physicochemical properties that collectively influence the three-dimensional conformation and binding interfaces. To achieve this, we adopt a graph-based representation. Structural interactions in epitope recognition depend strongly on physicochemical properties and tertiary chain organization; therefore, a graph-based representation is adopted to encode amino acid relationships more realistically. In addition, currently available datasets predominantly contain positive binding examples, whereas negative instances are commonly generated randomly and often lack biological meaning, which limits discriminative learning. Even in cases where real negative samples are available, such data alone may still be insufficient for effective training. To overcome this limitation without modifying model parameters, a tutorial-learning strategy is introduced to enrich the training data with informative synthetic negatives, facilitating a more robust learning process driven primarily by data rather than model complexity.

### Related work

Given the need for structure-aware prediction, it’s important to situate our work among existing TCR–epitope prediction approaches. The generation of T-cell receptor (TCR) diversity occurs through the random recombination of Variable (V), Diversity (D), and Joining (J) gene segments. Each V, D, and J segment encodes a distinct amino acid sequence, and their combination produces the TCR region responsible for antigen interaction (Ostmeyer et al., 2021). Consequently, each individual possesses a unique genetic immune repertoire. This inherent genetic and amino acid sequence diversity broadens the potential for immune response. This necessitates the development of predictive models capable of identifying correlations between TCRs and epitopes determined by V(D)J genes.

Prediction models like TCRex (Gielis et al., 2019) are available to immunology researchers through web applications. These epitope-TCR models, while differing in modeling approaches and feature sets (e.g., random forests, NLP techniques, and amino acid physicochemical properties (De Neuter N. et al., 2018)), share fundamentally similar classification approaches (Bi et al., 2019). Recent studies using such machine learning methods include: TCRex (Gielis et al., 2020), employing random forests and baseline prediction rates; TEPCAM, TITAN, and ATM-TCR (Chen et al., 2023; Weber et al., 2021; Cai et al., 2022), utilizing attention mechanisms; CLAIRE (Glazer et al., 2022), leveraging natural language processing (NLP); and the BERT-based TCRBERT (Wu et al., 2021; Jiang, Huo, & Cheng Li, 2023; Lu et al., 2021). Recently, a graph-based approach called HeteroTCR (Yu et al., 2024) has been proposed for TCR-epitope binding prediction. Unlike HeteroTCR, which primarily focuses on sequence-derived heterogeneous graph representations, our approach incorporates a broader set of physicochemical features and introduces a tutorial learning strategy for data augmentation. NetTCR-2.0 (Montemurro et al., 2021) uses a shallow convolutional neural network, and ERGO-II (Springer et al., 2021) applies a multilayer perceptron network with autoencoder-based encoding. TCR-Pred’s mathematical algorithm, MultiPASS, is based on the Bayesian method (Lagunin et al., 2000; Smirnov et al., 2023). Techniques such as meta-learning, contrastive learning, and ensemble learning have also been used to predict binding specificity (Fang et al., 2022; Gao, 2023; Xu et al., 2021). Each model requires training for a specific epitope or, in multi-class models, for a set of epitopes, necessitating sufficient training examples of TCRs with the corresponding epitope characteristic. These epitopes are not always readily available (Moris et al., 2021).

Epitopes and T-cell receptors (TCRs) are composed of amino acid sequences, each amino acid exhibiting unique structural properties. These variations are partially represented within prediction models. The Blocks Substitution Matrix (BLOSUM) is a substitution matrix employed to display relationships among the 20 canonical amino acids. Scores within the matrix, ranging from +11 to -4, quantify the degree of similarity between amino acids. In pairwise comparisons, positive scores indicate conservative substitutions, whereas negative scores represent radical substitutions. For instance, if an analysis is based on a 62% similarity threshold, this matrix is designated “BLOSUM62” (Henikoff & Henikoff, 1992). In the construction of this matrix, amino acid pairs with similar chemical properties (e.g., hydrophobicity or polarity) are assigned higher positive scores. Frequent amino acid substitutions (substitution probabilities) receive positive scores, and substitutions between amino acids with similar characteristics are considered conservative, thus receiving higher scores in the matrix. Conversely, substitutions between amino acids with differing properties are considered radical, often receiving lower or even negative scores (Henikoff & Henikoff, 1992). As similarity thresholds vary, the matrix itself undergoes corresponding changes. In applications such as TCR epitope binding prediction models, particularly in deep learning methods such as natural language processing (NLP) models, amino acid names necessitate conversion into numerical values. Consequently, BLOSUM matrices with varying similarity thresholds are frequently utilized for this transformation. For example, BLOSUM62 (Jurtz et al., 2018; Jokinen et al., 2019; Weber et al., 2021; Zhang et al., 2023; Wu et al., 2023; Essaghir et al., 2022), BLOSUM80 (Zhang et al., 2023), BLOSUM39 (Croce et al., 2024), and BLOSUM50 (Fischer et al., 2020) have been incorporated into the encoding process of prediction tools. In addition to BLOSUM, Simplified Molecular Input Line Entry System (SMILES), a notation employed to symbolize chemical formula representations of atoms, is also a tool used in model development (Weber et al., 2021).

The binding relationships of adaptive immune receptors (AIRs), including T-cell receptors (TCRs) and B-cell receptors (BCRs), have been extensively documented. Deep learning methodologies are frequently employed to predict both the binding reactivity and affinity of TCRs and BCRs (Yu et al., 2023; Isacchini et al., 2021). Yu et al. (2024) proposed an integrative approach utilizing amino acid sequences and three-dimensional (3D) structural information to predict antigen-binding reactions of adaptive immune receptors. A key finding of their study was the prediction of the specific protein chain region where binding occurs. The determination of the binding region is critically dependent on the 3D protein chain structure. In this context, 3D structural information derived from amino acid sequences, as provided by AlphaFold2, was utilized (Jumper et al., 2021).

In Figure 1a, the structure shown in green corresponds to Streptogrisin B (PDB: 2QA9), which consists of a total of 238 amino acids. In Figure 1b, the structure highlighted in orange represents Ovomucoid (QMYKY3), a short peptide composed of 56 amino acids. Figure 1b also depicts the binding process between the two molecules in three dimensions, where the regions associated with strong interactions are indicated. In the same figure, a molecular surface representation is provided to illustrate a possible three-dimensional appearance of the binding interface. During this interaction, contacts on the Streptogrisin B protease chain are predominantly observed around His at position 57, Ser at position 195, and Residue 119 (Protein Data Bank), while the binding region of QMYKY3 primarily involves amino acids between positions 19 and 21. Notably, in the larger structure 2QA9, the interacting residues are located at distant positions within the sequence (57, 119, 195), which suggests that methods relying solely on primary sequence information (e.g., language models) may be insufficient. Therefore, approaches accounting for secondary or tertiary structure—such as the graph-based modelling strategy adopted in this study—may provide a more reliable basis for binding prediction. The 3D conformation of 2QA9, shaped by its sequence and environmental conditions, forms a binding pocket into which the QMYKY3 peptide fits. The same principle holds for epitope recognition: a peptide such as CAAGPLLMATGT cannot present all of its amino acids on the binding surface simultaneously (Mariuzza et al., 2020; Szeto et al., 2020; Bradley, 2023). Each amino acid sequence adopts a unique 3D structure that is influenced by physicochemical properties including hydrophobicity, hydrophilicity, basic or acidic character, atomic volume, and mass. Jumper et al. (2021) highlighted that the chemical properties of amino-acid side chains, as well as their local interactions within binding regions, play a decisive role in determining the resulting 3D conformation. Moreover, the development of AlphaFold2 explicitly incorporated interaction information between different regions or domains within the same protein, further underscoring the importance of structural context (Jumper et al., 2021).

**Figure 1:**
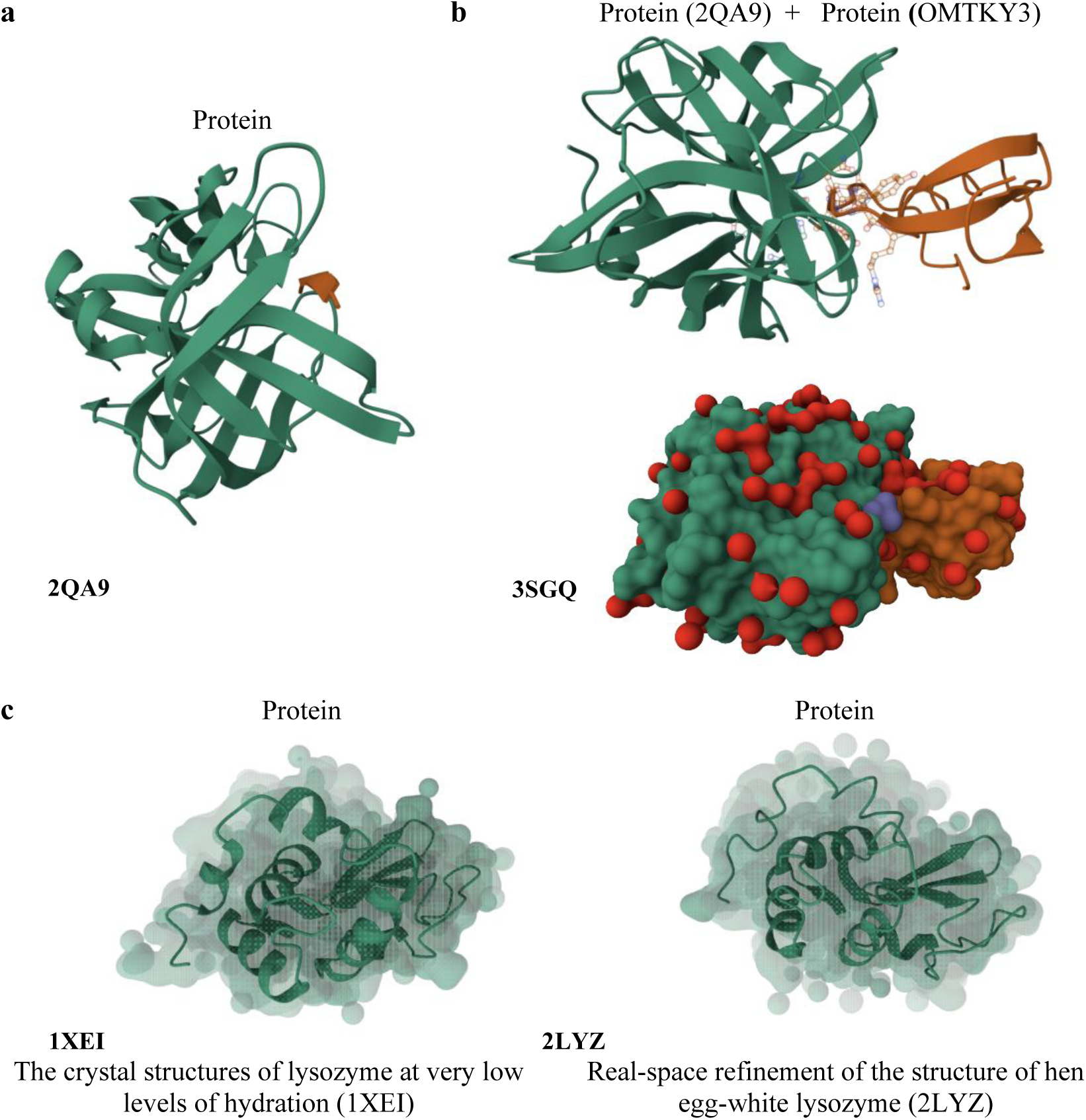
Simulation of protein–protein interaction and the effects of environmental conditions on protein structure (Protein Data Bank).

Due to their hydrophobic nature, some residues tend to avoid water and become buried inside the protein core, which may hinder intermolecular binding (Saotome et al., 2025). When the amount of water in the environment decreases, these hydrophobic amino acids move towards the surface, altering the overall 3D structure and consequently shifting potential binding regions. Thus changes in solvent composition modify the geometry and contact surface of the binding pocket and can dramatically alter binding propensity (Maurer & Oostenbrink, 2019). For example, in the lysozyme systems shown in Figure 1c, the chain architecture remains identical, and the only changing parameter is the amount of water present in the environment. Hydrophobic residues such as Isoleucine (I), Valine (V), Leucine (L), Phenylalanine (F), and Methionine (M) are more buried in the 2LYZ structure, thereby reducing the likelihood of binding. Consequently, environmental changes directly affect the shape of prospective binding pockets and constitute an essential determinant of molecular recognition. According to PeSTo predictions (pesto.epfl.ch; Krapp et al., 2023), protein binding is not predicted for 2LYZ, whereas DNA–RNA binding is estimated with a probability of 54 % for Gly at position 126. Similarly, in the case of 1XEI, the predicted protein–protein binding probability for Asn 113 and Arg 114 is 60 % and 53 %, respectively, while DNA–RNA interaction is not predicted for this structure (pesto.epfl.ch).

The example presented in Figure 1c summarizes only the case related to water parameters; however, numerous structural and environmental factors also play a role in protein binding. The interaction of proteins with other proteins or ligands is primarily governed by the intrinsic structural and dynamic properties of the protein itself. In particular, the amino acid sequence, secondary structure elements (e.g., α-helices and β-sheets), and overall three-dimensional folding determine the geometry, size, and chemical microenvironment of the binding site, thereby enabling a “lock-and-key”–like complementarity with ligands or partner proteins (Du et al., 2016; Bucher et al., 2011; Arai, 2018). The electrostatic surface potential, hydrophobic or hydrophilic patches, aromatic side chains, and other local chemical features directly influence the type and strength of interactions. Moreover, conformational flexibility and the ability to adopt multiple structural states facilitates binding-induced rearrangements via induced fit or conformational selection mechanisms (Du et al., 2016; Hancock, 2023; Liang et al., 1998). In addition, post-translational modifications such as phosphorylation, glycosylation, or disulfide-bond formation can remodel the molecular surface and reorganize the binding interface (Duan and Walther, 2015; Wang et al., 2022; Schofield et al., 2024). Oligomerization may generate novel interfacial binding regions and also promote cooperative effects, whereas internal cavities, tunnels, and specific water configurations influence both binding kinetics and energetic components (Maurer and Oostenbrink, 2019; Roberts and Mancera, 2008; Colloc’h et al., 2017; Meng et al., 2009; Li et al., 2022; Saotome et al., 2025). Furthermore, intrinsically disordered regions (IDPs/IDRs) may undergo folding upon binding, enabling interactions with multiple partners and rendering protein binding a highly dynamic and context-dependent process (Wong et al., 2020; Trivedi and Nagarajaram, 2022; Chen et al., 2022; Cermakova and Gibson, 2023). Beyond intrinsic determinants, several environmental conditions also markedly modulate binding, including temperature (Jahmidi-Azizi, 2021), pH and ionic strength (Shi et al., 2019), solvent polarity and water structure (Wernersson et al., 2022; Sree et al., 2025), metal ions and redox status (Majorek et al., 2024; Hartwig, 2013; Jevtović et al., 2023; Bromberg et al., 2022), exposure to light or UV radiation (Mondal, 2018), viscosity (Papaneophytou et al., 2014; Wernersson et al., 2022; Quaye et al., 2022), and macromolecular crowding (Köhn et al., 2021; Alfano et al., 2024). Consequently, any predictive framework aiming to estimate binding strength or specificity must incorporate multiple structural and environmental parameters simultaneously. Models (such as protein language model) that rely solely on a limited subset of inputs are unlikely to sufficiently capture the complex and multifactorial nature of biological binding events.

In the literature, a wide range of machine learning models have been employed that rely primarily on the protein’s primary sequence to address binding problems. Compared with primary-sequence-based CNN approaches, graph-convolutional models that integrate secondary or structural information have been reported to achieve substantially higher accuracy (Gligorijević et al., 2021; Lin et al., 2024; Jiao et al., 2023). Si and Yan (2024) emphasized that the use of Protein Language Models (PLMs) alone is not sufficient for reliable binding prediction. Their results show that inter-protein contact prediction clearly improves when PLMs outputs are combined with geometric graph representations. Similarly, Sagendorf et al. (2024) demonstrated that structural graph representations are highly effective for protein–nucleic-acid interactions, while Wu et al. (2023) showed that PLM-embedded graph models produce richer and more discriminative representations than sequence-only methods. Taken together, these studies indicate that multi-feature, structure-aware, graph-based models provide a clear advantage over classical transformer or language-model approaches that rely on fewer parameters and only primary sequence information.

This study introduces a prediction framework that departs from existing approaches in several notable ways. First, we propose a graph-based representation enriched with an extensive set of physicochemical features, enabling a substantially more informative encoding of amino acid characteristics than conventional sequence-based or low-feature models.While the current implementation does not directly incorporate three-dimensional coordinates, it captures key physicochemical properties that influence tertiary structure formation, serving as a practical proxy for structural relationships in the absence of extensive structural data. The relation-centric design of the graph structure also permits the systematic exploration of alternative representations by modifying nodes, node attributes, edges, and edge properties, thus providing flexible and scalable modelling capabilities. Second, we incorporate a tutorial learning strategy aimed at improving later training epochs. In particular, we address the well-known instability caused by negative samples in performance comparison studies, and we demonstrate that training with agent-generated negative data—absent in the original dataset—yields more robust outcomes. To our knowledge, this is the first study to enhance model performance solely through dataset manipulation without adjusting model parameters. Taken together, this methodology advances predictive modelling in computational immunology while also offering broader implications for data-centric approaches in machine learning.

The next section presents a clear problem definition, along with the model design and results for selected sample cases, illustrated with graphs. Section 3 highlights methodological distinctions within computational immunology research based on these results. Section 4 provides a discussion of the results. Finally, we detail the parameters and the preparations necessary for training and running the model.

## 2 PROBLEM DEFINITION AND RESULTS

This section defines the core problem underlying the study, outlines the methods developed to address it, and presents a comparative evaluation of the model performances.

### 2.1 Problem definition and model overview

We address the problem of predicting TCR-epitope binding interactions using a graph-based approach by representing the TCR and epitope sequences as structured feature-rich entities. Unlike methods that rely solely on primary sequence, our approach encodes residue-level physicochemical properties that collectively influence the three-dimensional conformation and binding interfaces. This enables the model to capture residue-level relationships and interaction patterns essential for binding recognition.

To illustrate the importance of structural context in TCR–peptide recognition, Figure 2 presents an example of the interaction between the influenza M1 GILGFVFTL epitope and an αβ TCR. Figure 2 a, shows how the influenza M1-derived GILGFVFTL peptide (residues 58–66) is presented by HLA-A*02:01 and recognized by a specific αβ T cell receptor. Contrary to typical TCR–peptide interactions that rely heavily on prominent side-chain contacts, GILGFVFTL exposes mainly backbone atoms, leading to an unusual docking geometry in which the TCR engages the peptide–MHC complex with a more diagonal orientation. This arrangement allows a conserved residue in the TCR β-chain CDR3 loop to insert into a well-defined notch above the peptide–MHC surface, forming the critical contact that underlies the immune dominant CD8⁺ T cell response to this epitope in HLA-A2⁺ individuals. Figure 2 b shows the 3D molecular structure following β2-microglobulin association. Figure 2 c depicts the determined interactions and their strengths among the GILGFVFTL peptide, the TCR α-chain, β-chain, and the antigen. Dark grey indicates strong binding, while light grey denotes weak binding. For example, glycine at position 4 of GILGFVFTL forms a strong contact with the TCR α-chain, whereas its interactions with the β-chain and antigen in that same region are comparatively weak (Stewart-Jones et al., 2003). This example underscores how considering 3D orientation and contacts can be crucial, hence our model encodes such structural relationships.

**Figure 2:**
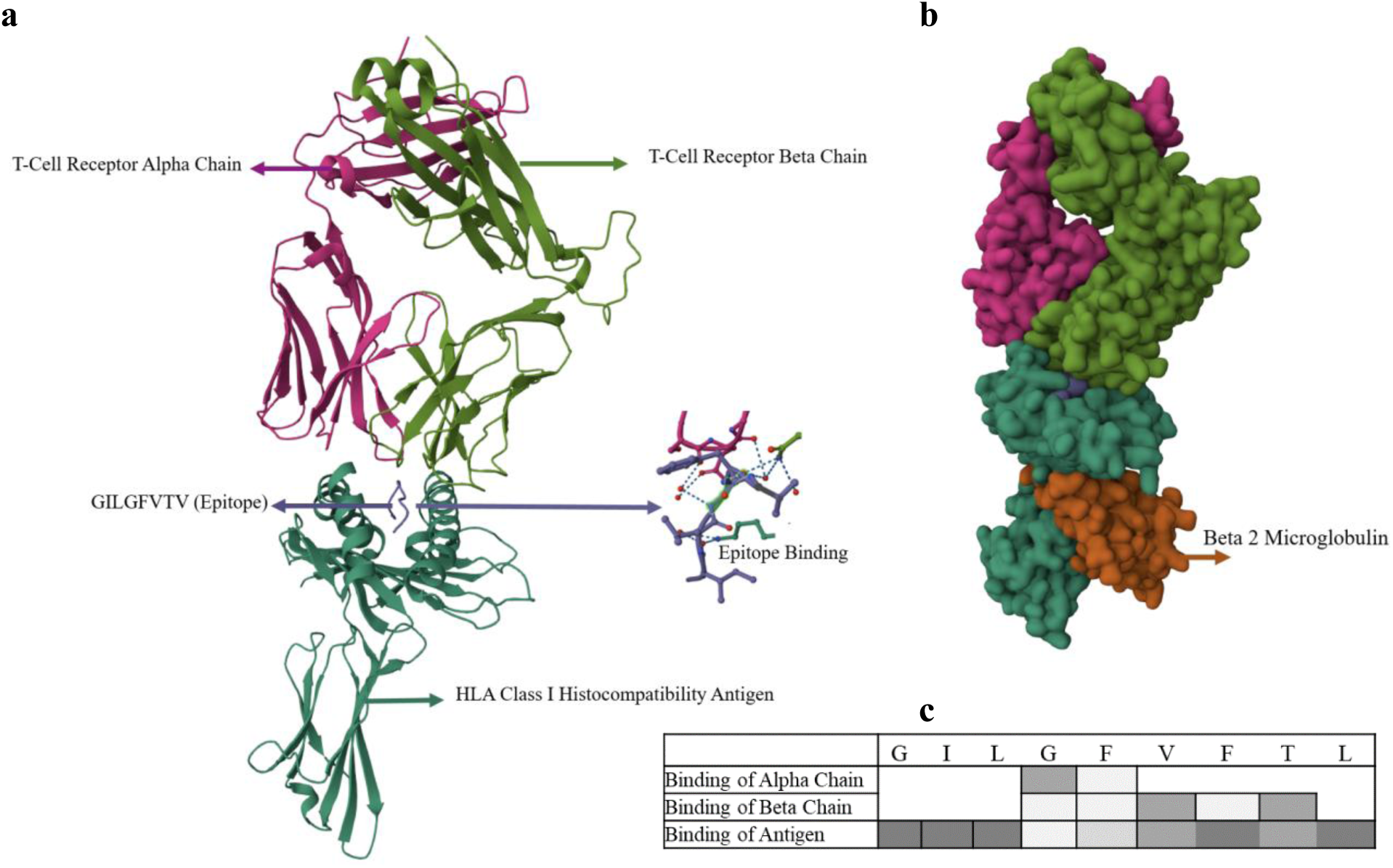
Interaction of the influenza M1 GILGFVFTL epitope with HLA-A*02:01 and the cognate αβ TCR (Protein Data Bank).

Laboratory results provide the basis for generating datasets essential to predictive model development. Figure 2, schematises the processing of peptides in the antigen production process. This environment utilizes various substances and cytokines to induce dendritic cell maturation and activation. Specific enzymes and auxiliary molecules (e.g., Proteasome Complex, TAP, ERAP) are employed to investigate MHC I and MHC II molecule processing and presentation (Evnouchidou and Endert, 2019; Saveanu et al., 2005). Following proteolysis, resulting protein fragments facilitate antigen presentation on both major histocompatibility complex (MHC) class I and class II molecules. These fragments, termed peptides, consist of amino acid sequences presented on these MHC molecules. Discrete regions within these amino acid sequences are designated epitopes. The interaction, or binding, of these epitopes with T-cell receptors (TCRs) expressed on T-cells initiates immune system activation, subsequently inducing antibody production. Identification of specific epitope-TCR binding pairs is a critical resource for therapeutic drug and vaccine development. However, challenges arise from substantial viral structural diversity, extensive genetic variability, and ongoing mutational processes, yielding a broad spectrum of epitope-TCR binding instances. Conventional laboratory methodologies are frequently subject to limitations including protracted timelines, substantial financial expenditures, and constraints in available resources such as equipment, raw materials, and personnel. These limitations impede efficient development of effective therapeutic interventions. Consequently, a machine learning model capable of accurately predicting potential epitope-TCR pairings based on existing data would provide substantial support for diverse therapeutic development efforts.

### 2.2 Graph Model for better generalization and Tutorial Learning

The details of the multi-feature graph model and the agent-based negative data generation for tutorial learning are presented below.

#### 2.2.1 Bipartite Graph Representation of TCR Sequences

We represent each TCR–epitope pair as a bipartite graph, where one partition corresponds to the amino acids of the TCR and the other to the ordered positions of the epitope. For each epitope, the constructed graphs incorporate multiple biochemical features assigned at the TCR nodes. In graph theory, networks can be classified as homogeneous or heterogeneous based on the uniformity of their node types. Homogeneous networks consist of nodes of a single type (e.g., individuals, stations, or routers). Within this classification, a homogeneous graph is equivalent to a simple graph, as opposed to a bipartite graph. Formally, a homogeneous (or simple) graph, denoted as *G* = (*U*, *E*), is defined by a set of nodes, *U*, and a set of edges, *E*, that connect these nodes. An edge connecting a node *u* ∈ *U* to a node *v* ∈ *U* is represented as (*u*, *v*) ∈ *E* (Giamphy et al., 2023). In contrast to such homogeneous structures, the bipartite formulation used in this study assigns TCR residues and order positions to two distinct node sets, enabling a more structured representation of secondary and three-dimensional features.

A bipartite graph, denoted as *G* = (*U*, *V*, *E*), consists of two disjoint sets of nodes, *U* and *V*, representing distinct domains, and a set of edges, *E* ⊂ *U* × *V*, connecting these nodes. A defining characteristic of bipartite graphs is the restriction of edges to connections between nodes in the distinct sets U and V; intra-set connections are disallowed. Consequently, such graphs are classified as heterogeneous graphs. An illustrative example is presented in Figure 3, clearly demonstrating the two distinct sets and their interconnectivity. Within this framework, set *U* can be interpreted as representing amino acid nodes, while set *V* represents sequence nodes. Let *u_i_* represent the ith node in *U* (where *i* = 1,2, …, *n*), and *v_j_* represent the jth node in *V* (where *j* = 1,2, …, *m*). An edge (*u_i_*, *v_j_*) signifies the presence of a connection between the ith node *u_i_* in *U* and the jth node *v_j_* in *V*. Given *n* = |*U*| and *m* = |*V*|, an *n* × *m* adjacency matrix is generated. In Figure 3, the matrix W represents the adjacency matrix between the node sets *U* and *V*. Specifically, *w_ij_* = 0 indicates the absence of a connection between the ith node in *U* and the jth node in *V*, while *w_ij_* = 1 indicates the presence of such a connection. It is important to note that in this graph, edges are unweighted; all connections are assumed to have uniform strength, represented by a value of 1. Furthermore, no attribute set has been assigned to the edges in this study. However, a comprehensive set of features (attributes) has been defined for the nodes.

**Figure 3:**
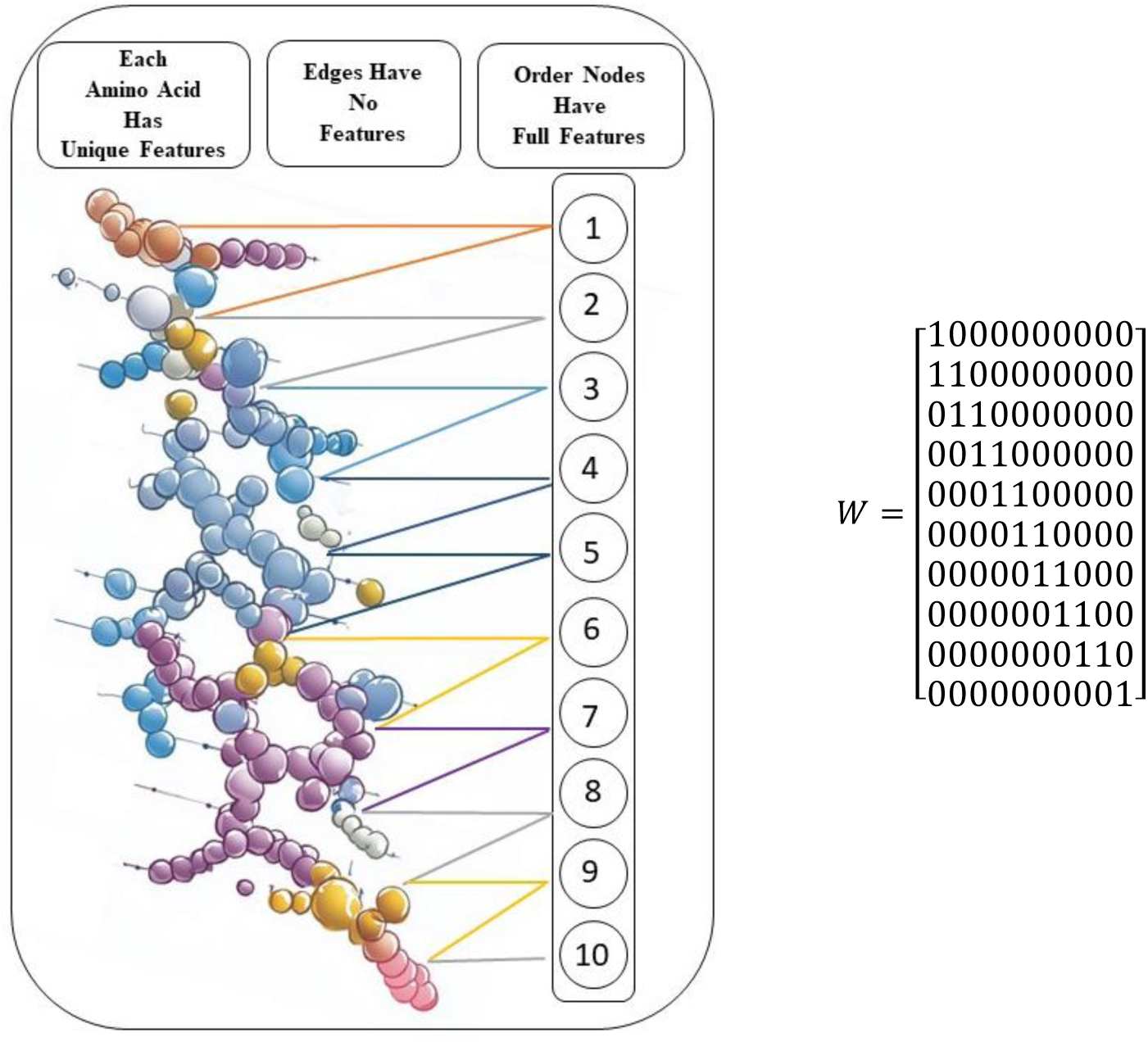
Construction of a bipartite graph according to the amino acid properties using the amino acid sequence of the TCR.

Following the design of the amino acid sequence as a bipartite graph, a graph-based machine learning model was selected for implementation. Graph Neural Networks (GNNs) are popular in the domain of graph representation learning (Li et al., 2016; Hamilton et al., 2017; Kipf & Welling, 2017; Velickovic et al., 2018; Xu et al., 2018). GNNs have been successfully integrated with established deep learning models and adapted to address contemporary challenges, including anomaly detection and image processing (Kong et al., 2024; Dhadhania et al., 2024; Yu et al., 2023). These models commonly employ a recursive neighborhood aggregation, or message-passing paradigm, wherein each node updates its feature vector by aggregating the feature vectors of its neighboring nodes (Xu et al., 2018; Gilmer et al., 2017).

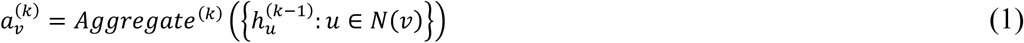

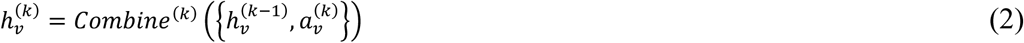

The aggregate function combines information from a node’s neighbors. For a given node u, the feature vectors of its neighbors (*N*(*v*)), represented as 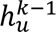, are collected and combined by an aggregation function (*Aggregate_k_*). This process generates a summary of the neighbors’ features, denoted as 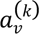. Thus, 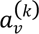 represents the neighbors’ influence on node *v* after *k* iterations (Xu et al., 2018).

The *Combine* function merges a node’s features with a summary of its neighbors. The updated node feature vector, denoted as 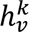, is generated by combining the node’s prior feature vector, 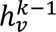, with the summarized attribute vector of its neighbors, *a*^(*k*)^. Here, 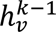 represents node *v*’s feature vector at the (*k*-1)th iteration, and *N*(*v*) represents the set of neighbors of node *v*. This approach allows Graph Neural Networks (GNNs) to update a node’s features by incorporating information from its neighbors. Consequently, the node’s feature vector learns a broader context at each iteration. After *k* iterations, all node features are determined.

In the second stage, the classification of a graph with known node features is determined. This process employs a Readout function, as detailed in

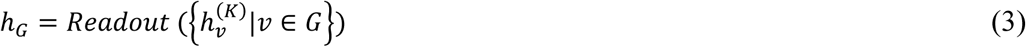

The Readout function, denoted as ℎ*_G_*, generates a global graph representation by aggregating the final feature vectors of all nodes. This aggregation may be performed through summation, averaging, or maximization of the values across all nodes. The value ℎ*_G_*, obtained via Equation 3, is then assigned to a class using a function defined as *y_G_* = *f*(ℎ*_G_*). For example, a comparison against a specific threshold can differentiate labels between graphs. Functions commonly utilized in machine learning, including softmax, sigmoid, and linear functions, are frequently employed for *f*(ℎ*_G_*) (Xu et al., 2018).

The design illustrated in Figure 3 comprises two distinct components. A 3D representation depicts nodes corresponding to amino acids. A sequence representation indicates the order of each node. Consequently, a bipartite graph is formed, with one component representing amino acids and the other representing order. The bipartite graph consists of two disjoint node sets: Position Nodes (Set V, nodes 1-20) representing sequential positions and Amino Acid Identity Nodes (Set U, nodes 21-40) representing the 20 standard amino acids. An edge connects position i to amino acid node j if amino acid j appears at position i of the TCR sequence. This bipartite formulation allows the model to learn position-amino acid associations while incorporating physicochemical features as node attributes. There are 20 distinct amino acids, including Alanine (Ala), Cysteine (Cys), Aspartic Acid (Asp), Glutamic Acid (Glu), Phenylalanine (Phe), Glycine (Gly), Histidine (His), Isoleucine (Ile), Lysine (Lys), Leucine (Leu), Methionine (Met), Asparagine (Asn), Proline (Pro), Glutamine (Gln), Arginine (Arg), Serine (Ser), Threonine (Thr), Valine (Val), Tryptophan (Trp), and Tyrosine (Tyr). These amino acids are encoded through numerical assignment, with numbers 21 through 40 assigned accordingly. Each number uniquely identifies an amino acid (e.g., 21 for Alanine (Ala, A), 22 for Cysteine (Cys, C), and 40 for Tyrosine (Tyr, Y)). Each amino acid possesses specific characteristics. Amino acids exhibit diverse physiochemical properties, including variations in hydrophobicity, hydrophilicity, essentiality, basicity, acidity, volume, and mass. Furthermore, their behavior is influenced by factors such as pH and side chain structure. As detailed on our public repository^1^, 47 different features were assigned in binary form, indicating whether a given feature is present in an amino acid. While the current implementation does not directly incorporate three-dimensional coordinates, this comprehensive physicochemical feature set serves as a practical and computationally tractable proxy for structural context. All features were fully assigned to the first 20 nodes (order nodes) within the model. Separate experiments were conducted with no feature assignment and full feature assignment to these virtual nodes. Full feature mapping resulted in improved performance. This design is shown schematically in Figure 3.

#### 2.2.2. Tutorial Learning

##### Algorithm 1

**Tutorial Learning Algorithm**

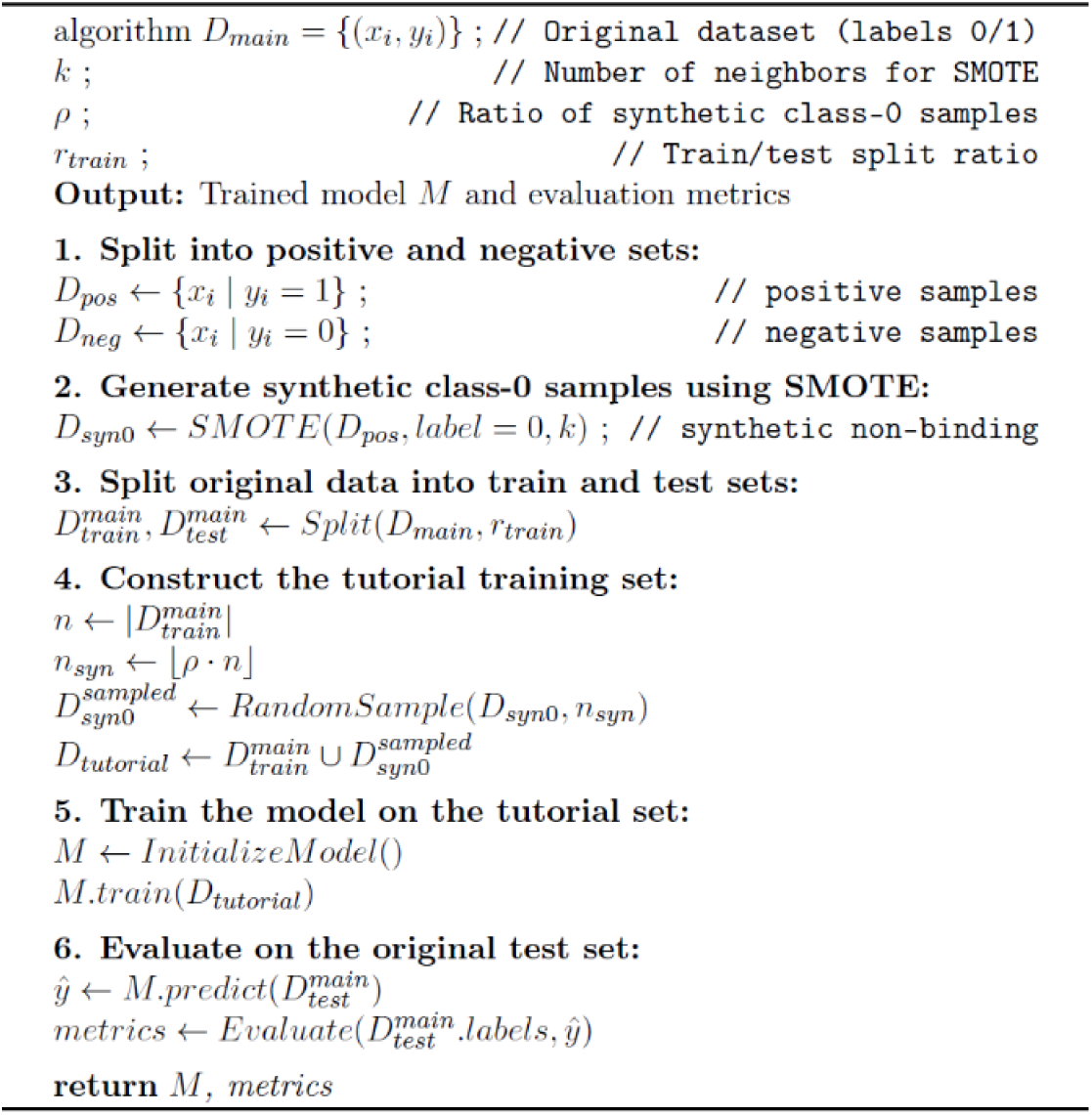

The key insight is that SMOTE operates in the feature space of the graph embeddings, generating synthetic negatives that lie in the vicinity of positive samples, thereby creating ‘tutorial’ examples that guide the model toward more discriminative decision boundaries.

This algorithm applies the Tutorial Learning approach shown in Figure 4. In the first step, the original dataset *D_main_* is divided into positive samples *D_pos_* and negative samples *D_neg_*. In the second step, a neighborhood value (*k*) for the SMOTE algorithm and the ratio of synthetic negative samples (*ρ*) are defined. SMOTE generates synthetic samples by interpolating between a selected instance and its k-nearest neighbors in the feature space (Chawla et al., 2002). In this study, SMOTE is applied to the flattened feature representation of the graph, where each TCR-epitope pair is vectorized by concatenating node features and adjacency matrix. Specifically, we used *k=5* for nearest neighbor selection and generated synthetic negative samples at a ratio of 1:1 relative to the real negative samples in the training set. This approach generates new synthetic negatives that lie in the vicinity of the positive samples, effectively acting as tutorial examples to guide the model toward more discriminative decision boundaries.

**Figure 4:**
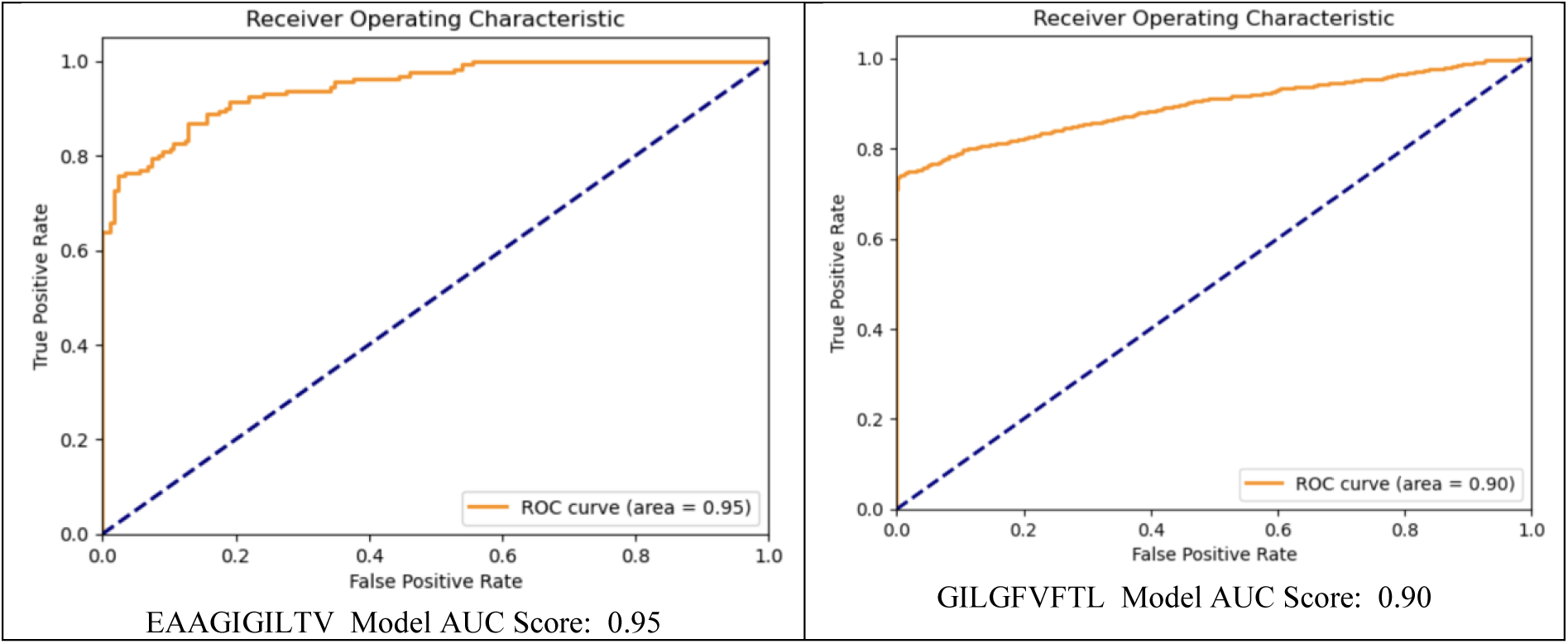
Pseudo code of Tutorial learning Algorithm.

In summary, based on these parameters, synthetic negative samples *D_syn_*_0_ are generated from the neighborhood of the positive samples in order to strengthen the model’s ability to distinguish between the classes. In the third step, *D_main_* is split into a training set *D^main^* and a test set *D^main^*. In the fourth step, a subset of the synthetic samples *D_syn_*_0_ is combined with the training set to form the tutorial training set. The purpose of this step is that the synthetic negatives act as “tutorial examples” and help the model to better recognize real negative samples. In the fifth step, the model is trained on the extended tutorial training set (*D^main^* + *D_syn_*_0_). In the final step, the trained model is evaluated using the test set *D^main^*, which contains only real samples. In this way, the contribution of the synthetic samples to the overall classification performance can be objectively assessed.

### 2.3 Detailed analysis of predictive power of individual components in the model

Modeling studies, based on the aforementioned structural description, were conducted for five frequently reported epitopes.

The performance of the proposed model is presented in Figure 5 and compared with the results in Figure 2. The model achieved an AUC of 0.95 for the EAAGIGILTV epitope, as illustrated in the corresponding graph. For the GILGFVFTL epitope, the AUC was 0.90. The AUCs for the YVLDHLIVV, KLGGALQAK, and KAYNVTQAF epitopes were 0.89, 0.71, and 0.68, respectively. These five epitopes are well-established in the literature and have associated coding resources from prior studies. The availability of these prior binding and non-binding datasets facilitated accurate comparisons with existing research.

**Figure 5:**
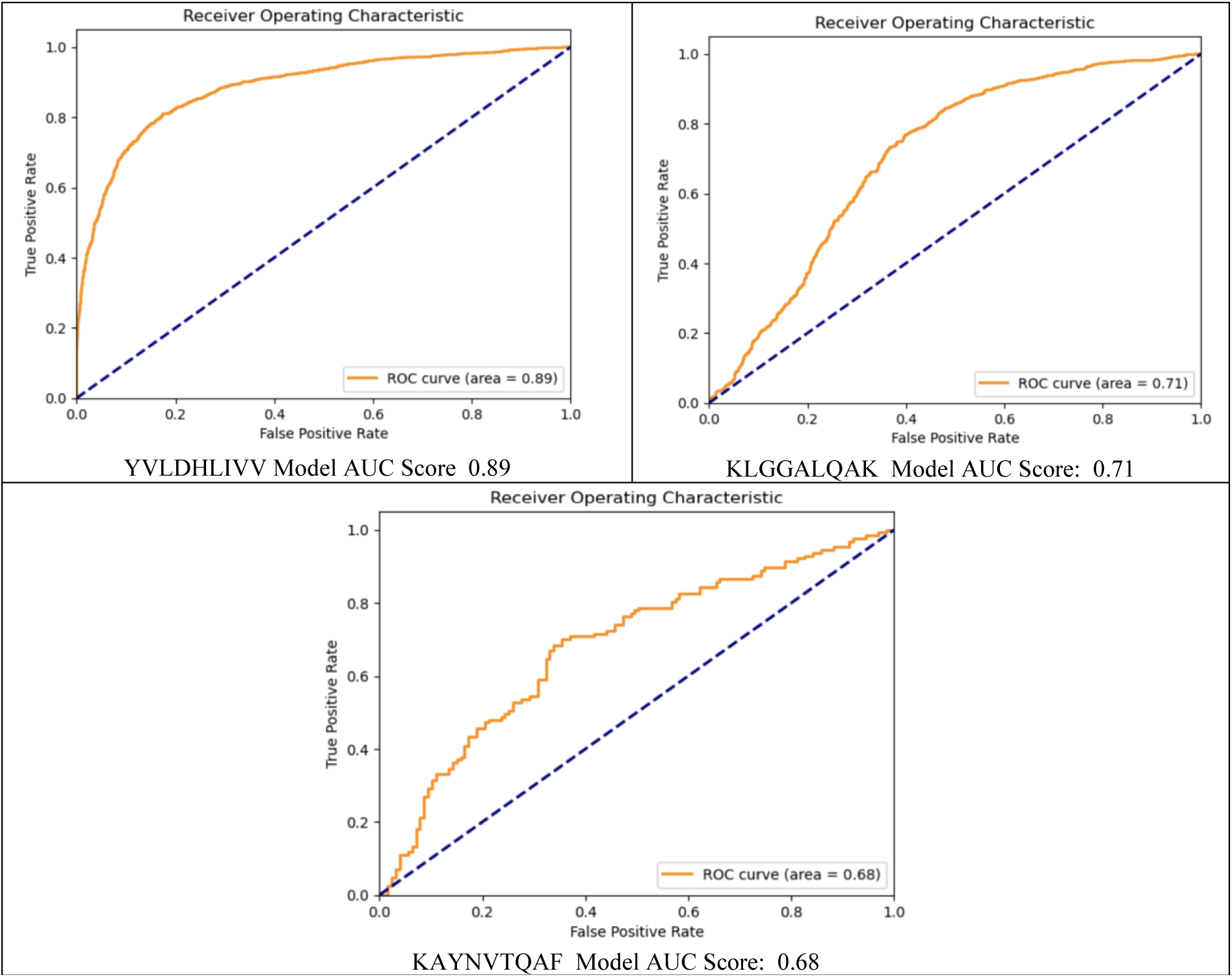
Model performance outputs for different epitopes using the ROC curve.

Table 1 compares models predicting binding and non-binding interactions. The datasets used in these studies are available on their corresponding GitHub repositories. Each study performed a performance comparison using its specific dataset. However, most studies in the literature independently generate their non-binding (0) data, while binding (1) data are typically downloaded from databases such as VDJdb, McPass, and IEDB. Consequently, model comparisons are conducted using datasets with at least 50% data discrepancy due to variations in the non-binding dataset. This indicates that the selection of non-binding data significantly influences model performance. Section 5.3 demonstrates this variation using a dataset generated with the Synthetic Minority Oversampling Technique (SMOTE) (Chawla et al., 2002). While research studies often describe the model architecture, details such as learning rate, batch size, number of hidden layers, seed value, dropout rate, optimization method, train-test split ratio, activation function, and alpha-beta ratio are generally not disclosed. Consequently, the performance of these undisclosed parameters on different datasets remains uncertain. The marginal performance differences observed in most comparisons emphasize the importance of these details.

**Table 1:** Performance differences between TCR-epitope machine learning models.

| Epitope-TCR Models | Node Feature | Dataset Samples | AUC | Accuracy | Precision | Recall | F1 |
|---|---|---|---|---|---|---|---|
| **EAAGIGILTV- Graph Desing** | 35 | 968 | 0.95 | **0.86± 0.02** | 0.84 | 0.87 | 0.85 |
| EAAGIGILTV - TCR-Pred - (Croce, et al., 2024) | BLOSUM89 | 968 | - | 0.84 | - | - | - |
| EAAGIGILTV - TCR-Pred - (Smirnov. et al., 2023) | - | 968 | - | 0.84 | - | - |  |
| **GILGFVTL - Graph Desing** | 44 | 10336 | **0.90** | 0.85± **0.02** | 0.94 | 0.76 | 0.84 |
| GILGFVTL - ATM-TCR-(Cai. et. al., 2022) | BLUSUM | 10336 | 0.88 | - | - | - | - |
| **YVLDHIVV - Graph Desing** | 322 | 16126 | **0.89** | 0.82± **0.02** | 0.82 | 0.79 | 0.80 |
| YVLDHIVV - ATM-TCR-(Cai. et. al., 2022) | BLOSUM | 16126 | 0.88 | - | - | - | - |
| **KLGGALQAK - Graph Desing (Tutorial)** | 886 | 5500 | **0.71** | 0.67± **0.02** | 0.62 | 0.89 | 0.72 |
| KLGGALQAK- ATM-TCR-(Cai. et. al., 2022) | BLOSUM | 25930 | 0.66 | - | - | - | - |
| **KAYNTVQAF - Graph Desing (Tutorial)** | 886 | 1604 | **0.68** | **0.67± 0.02** | 0.67 | **0.68** | **0.67** |
| KAYNTVQAF-TEPCAM- (Chen et. al., 2024) | - | 1209 | 0.65 | 0.60 | - | 0.62 | 0.61 |

A direct one-to-one performance comparison across methods is challenging because each study employs different negative sample generation strategies and train-test splits. Table 1 presents reported values alongside our results for reference, with the understanding that experimental conditions vary. Therefore, performance values presented in the referenced articles were compared using the authors’ original datasets (encompassing both binding and non-binding data). A further limitation encountered was the difficulty of replicating published results due to subsequent updates in certain code libraries, which precluded performance comparisons using alternative datasets. Each study’s results, as presented in Table 1, were derived from its respective dataset. Each study’s results, as presented in Table 1, were derived from its respective dataset. We optimized a separate graph model for each epitope dataset, adjusting model complexity to match sequence complexity (hence the feature counts differ, see Table 1). Two challenging epitopes (KLGGALQAK and KAYNTVQAF) also employed the tutorial learning strategy to generate additional negative samples. Consequently, some performance values in Table 1 are absent, as they were not disclosed in certain publications. The italic entries in Table 1 indicate the results of this study and highlight its performance advantages.

Table 1 displays the “ Node Feature “ column, indicating the quantity of features employed in the epitope models proposed within this study. The values highlighted in italics in Table 1 indicate the superior performance achieved in this study, and the corresponding row represents the models employed in our approach. The remaining rows correspond to other high-performing studies in the literature whose datasets (positive and negative dataset) are publicly available. While the features represented by BLOSUM entail a combined comparison, they fundamentally constitute a single feature, previously discussed in this document. For each TCR-epitope match, distinct model designs were generated, resulting in variations in the number of features utilized. In the “ Dataset Samples “ column, the “Tutorial Learning” method was implemented due to the complex structures of the KLGGALQAK and KAYNTVQAF models. This implementation led to variations in the sample size. Comprehensive details regarding the “Tutorial Learning” method are provided in Section 5.3 of this study.

Analysis of referenced studies reveals distinct methodologies for TCR-epitope match prediction. Croce et al. (2024) employed the BLOSUM similarity ratio as a feature, reporting a value of 0.89, with observed variation across matches. Their work introduced MixTCRpred, a Transformer Encoder-based natural language processing model achieving 84% accuracy for the EAAGIGILTV epitope. Smirnov et al. (2023) utilized a 50% proportionally labeled dataset, converting molecular structure formulas into matrices and applying a naive Bayes classifier. This resulted in MultiPASS, a binary prediction model also achieving 84% accuracy. These studies demonstrate disparate yet accurate approaches to TCR-epitope match prediction.

In this study, data previously used by Cai et al. (2022) for GILGFVTL, YVLDHIVV, and KLGGALQAK was also employed. Utilizing a Multi-Head Self-Attention Model, Cai et al. reported AUC-ROC scores of 88%, 88%, and 68% for these respective datasets. Comparatively, this study achieved scores of 90%, 89%, and 71% using the same datasets. A detailed discussion regarding the prediction of KLGGALQAK is provided in Section 5.3. Chen et al. (2024) proposed the TEPCAM model for epitopes in contexts with limited datasets. Using the same dataset for the KAYNTVQAF epitope, the area under the receiver operating characteristic curve (AUC) was improved from 65% to 68%. This performance improvement was achieved by adopting the “Tutorial Learning” approach described in Section 5.3. Additionally, because each epitope required a separately tailored model, our findings further underscore the need for a more generalized solution—a point elaborated in the limitation discussion in Section 3.1.

## 3 DISCUSSION

The proposed model incorporates amino acid properties to enhance prediction accuracy. Comparison with state-of-the-art methods underscores the importance of realistic dataset generation and feature-rich graph representations. Challenges include dataset variability and the high computational cost of graph-based models. Our results confirm that a multi-feature, structure-aware approach yields higher accuracy than single-feature models, validating the benefit of incorporating diverse amino acid properties. The potential to determine which feature is more influential in possible TCR-epitope matches is significant for laboratory studies. For instance, acidity may be critical in some matches, while hydrophobicity may be more important in others. This information provides a valuable guide for establishing optimal laboratory conditions.

In cases where Tutorial Learning was applied, we observed improved prediction performance solely through the inclusion of synthetic negative samples, without modifying the underlying model. This approach is particularly valuable for structurally complex epitopes where learning is inherently challenging. While conventional model-parameter optimization can improve predictive performance only up to a certain level, the additional training enabled by Tutorial Learning provides an extra performance gain. The number of case studies was limited to five due to the scarcity of publicly available datasets that both report high prediction performance and provide accessible benchmark data. Moreover, in this study prediction was performed using multiple models that were not entirely identical. Although the fundamental approach relied on a bipartite graph structure, the model parameters (e.g., number of layers, learning rate) varied across models. This variation mainly results from the different levels of structural complexity observed in protein chains; more complex chains required a larger number of parameters. This limitation may be alleviated in future work. If a larger number of cases are available, a pool of pretrained models could be created, after which a hyper-model could be designed. In the future, a library of pre-trained models could be developed, so that for any new epitope one can select the closest matching model from the library instead of training a new model from scratch. Since this approach bypasses an additional training stage, it would allow evaluation based solely on test performance for epitopes with limited experimental observations. Thus, high-performing models could still be employed to make predictions even for rare epitopes.

A key problem highlighted in this study is the lack of standardized inclusion of unbinding (negative) data across training datasets, which complicates fair performance comparisons between models in literature. Consequently, it may be possible to claim superior performance by using less robust models combined with more discriminative or artificially curated unbinding samples.

This study emphasizes that data repositories containing binding interactions should also systematically include experimentally validated unbinding datasets. Such data are critically important: while artificially generated datasets may exhibit high separability, laboratory derived data often reveal that even a single amino acid variation can abolish binding, underscoring the subtlety and complexity of these interactions. Therefore, unbinding data are at least as important as binding data. Moreover, environmental conditions such as temperature, pressure, and ionic context should be reported for both binding and non-binding cases. Incorporating these factors would enable the development of more realistic and reliable predictive models.

In this study, for TCR–epitope interactions, epitope-specific architectures are proposed by tuning distinct parameters for each epitope. Although employing a single, unified model may appear more practical, it has several limitations. For instance, a model may achieve high overall performance by accurately predicting relatively simple and well represented binding cases, while failing to generalize to less frequent or more complex interactions that remain underexplored.

As a result, despite high aggregate performance metrics, such models may not effectively capture rare or specialized binding patterns. Therefore, this study argues that case-specific (epitope-tailored) models are more suitable, particularly for interactions with limited sample sizes and higher biological complexity.

### 3.1 Limitation

Comparisons are based on reported values from original publications. Due to differences in negative sample generation and train-test splits, these comparisons should be interpreted as indicative rather than strictly controlled.

One limitation of the current model is that it does not consider certain environmental factors. For instance, while the scenario focuses exclusively on TCR–epitope binding, other molecules might pre-occupy the relevant binding pocket, as illustrated in Figure 2, which could lead to incorrect predictions. Beyond this study, this issue represents a broader limitation in the field. Binding is not solely determined by the interaction between two molecular structures; it is also significantly influenced by environmental conditions such as ion concentration, water content, temperature, and hormonal factors. Therefore, it is crucial that binding data in existing databases be augmented with contextual environmental information to enable more accurate and biologically meaningful analyses.

Another improvement suggestion is schematized in Figure 6. The model presented above in Figure 6 represents an enhanced version of the design proposed in this study. However, there are several reasons why this improved model could not be fully applied. First, effective training of the developed graph model requires a large diversity of T-cell and epitopes positive samples. In addition, in this work the protein chain length was restricted to 20 amino acids. Extending the chain length would require integrating both environmental and secondary chain features into the model, which would substantially increase the computational cost of training a two-variable structure (TCR and epitope). Therefore, both the limited dataset and the high computational demand constitute major constraints of this study. Under the current conditions, the training time for the five models implemented in this work ranged between 20 and 35 minutes using a computer with a 12th Gen Intel Core i9-12900H (2.50 GHz) processor and 32 GB RAM. Additionally, while we report single-run performance with fixed seeds, the consistent improvement across multiple epitopes suggests robustness. Future work will incorporate multi-seed validation to provide statistical confidence intervals.

**Figure 6:**
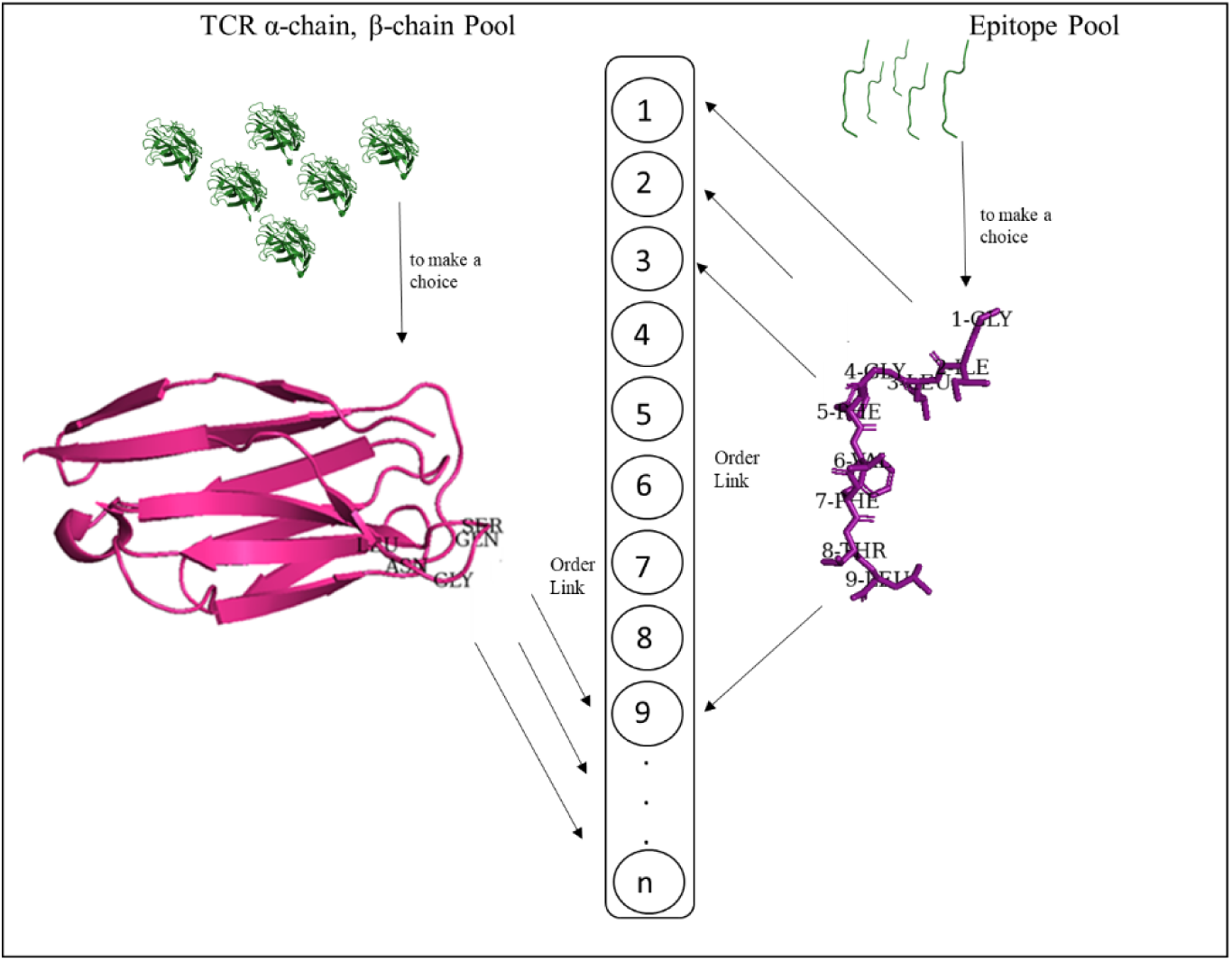
Enhanced graph model design.

## 4 CONCLUSION

This study introduces a novel bipartite graph model for predicting T-cell receptor (TCR)-epitope binding, demonstrating substantial performance gains across datasets of diverse epitope characteristics. Despite relatively small sample sizes, the EAAGIGILTV and KAYNTVQAF epitopes exhibited varying performance outcomes, underscoring the challenge of achieving robust predictions across diverse epitope types using a single model with fixed parameters. This challenge is primarily attributable to the distinct chemical compositions of epitopes and TCRs. Amino acid properties, such as hydrophobicity, hydrophilicity, essentiality, basicity, acidity, volume, and mass, influence both their three-dimensional structures and corresponding binding surfaces. This study hypothesizes that a more accurate simulation of these properties in model design will enhance binding prediction accuracy. Consequently, the model incorporates diverse features beyond a single network binding. However, feature set effectiveness varies across potential epitope-TCR bindings, resulting in variable feature utilization depending on the potential binding (epitope) type. Despite large sample sizes, the models for KLGGALQAK and KAYNTVQAF epitopes exhibited suboptimal learning performance, potentially due to non-binding dataset diversity or binding data chemical structure variations. These datasets also presented a significant overfitting risk. Conversely, a more balanced dataset for the YVLDHIVV epitope, with 75% allocated for training, yielded notably high performance. These findings demonstrate the unique characteristics of each epitope model. Future research will investigate multi-epitope scenarios and integrate additional physicochemical features to further refine predictions.

### 4.1 Future Work

To address the current epitope-specific limitation, future work will explore transfer learning and meta-learning approaches to develop a unified model capable of generalizing across epitopes.

## 5 MATERIALS AND METHODS

### 5.1 Data curation

Available databases (VDJdb, McPass, IEDB) exclusively contain positive (binding) results, indicative of interactions between epitopes and peptides. While this is a conventional approach, diverse strategies are employed without generating corresponding negative data. Consequently, studies have employed randomized datasets. For example, Moris et al. (2021) generated negative samples utilizing the interaction map recognition method. This method is predicated on the recognition of epitope sets facilitated by the polymorphic nature of MHC genes and the quasi-random somatic V(D)J recombination of TCRs.

Cai et al. (2022) constructed ATM-TCR models using binding pairs sourced from three databases: VDJdb, McPass, and IEDB. Non-binding pairs were generated via random recombination of existing positive pairs. The resultant TCRβ CDR3-epitope pairs formed the training dataset. Consequently, the β chain was also employed in the present study.

In the TEPCAM model, Chen et al. (2024) established a TCRβ CDR3-epitope dataset derived from the same databases, implementing an experimental approach (Gielis et al., 2019) to mitigate randomness within the non-binding dataset. For non-binding data generation, 250,000 unique chains were acquired from 587 healthy individuals, from which 100,000 were randomly selected. As documented in the limitations section of Chen et al. (2024), this dataset is confined to two viruses (yellow fever virus (YFV) and cytomegalovirus (CMV)) and a limited cohort of 587 individuals. It is posited that a modification in the virus type would likely engender the formation of disparate non-binding pairs, consequently necessitating the development of a novel model for machine learning applications.

To generate the 10X dataset, T cells were subjected to a panel of 50 peptide-MHC multimers via the 10X assay method. A negative TCR was defined as a TCR exhibiting no binding affinity to any of the tested peptides (Montemurro et al., 2022). Utilizing the negative dataset from Montemurro et al. (2022), an accuracy of 0.836 was attained for EAAGIGILTV (Croce et al., 2024). Consequently, dataset standardization remains incomplete. Therefore, for the purpose of ensuring accurate comparisons in this study, established datasets were employed. In addition, the datasets used in our study are shared in the acknowledgment section of the “doi” and “github” pages of the related studies.

### 5.2 Details of the model architecture

The following section describes the GNN models developed for five distinct epitope-specific settings. Although these may appear as five separate models, they are in fact parameterizations of the same bipartite architecture illustrated in Figure 3, adapted with epitope-specific configurations. The base model is built upon a single attention mechanism; however, due to the complex and binding-specific features of epitopes such as solvent interactions (e.g., water and ions), polarity, and side-chain structures the models exhibit variations in the number of layers, learning rate, batch size, and the choice of specialized deep learning components. The primary reason for these variations lies in the intrinsic chemical diversity of each epitope, which necessitates tailored modeling strategies to effectively capture their distinct biochemical properties.

The model constructed for EAAGIGITV, depicted in Figure 7, employs a less complex architecture, incorporating 100 hidden layers. This model was trained using 968 data points, with 65% (315 binding and 315 nonbinding) allocated for training. Details regarding the models developed for GILGFVTL, YVLDHIVV, KLGGALQAK, and KAYNTVQAF are provided in Figures 8, 9, 10, and 11, respectively.

**Figure 7:**
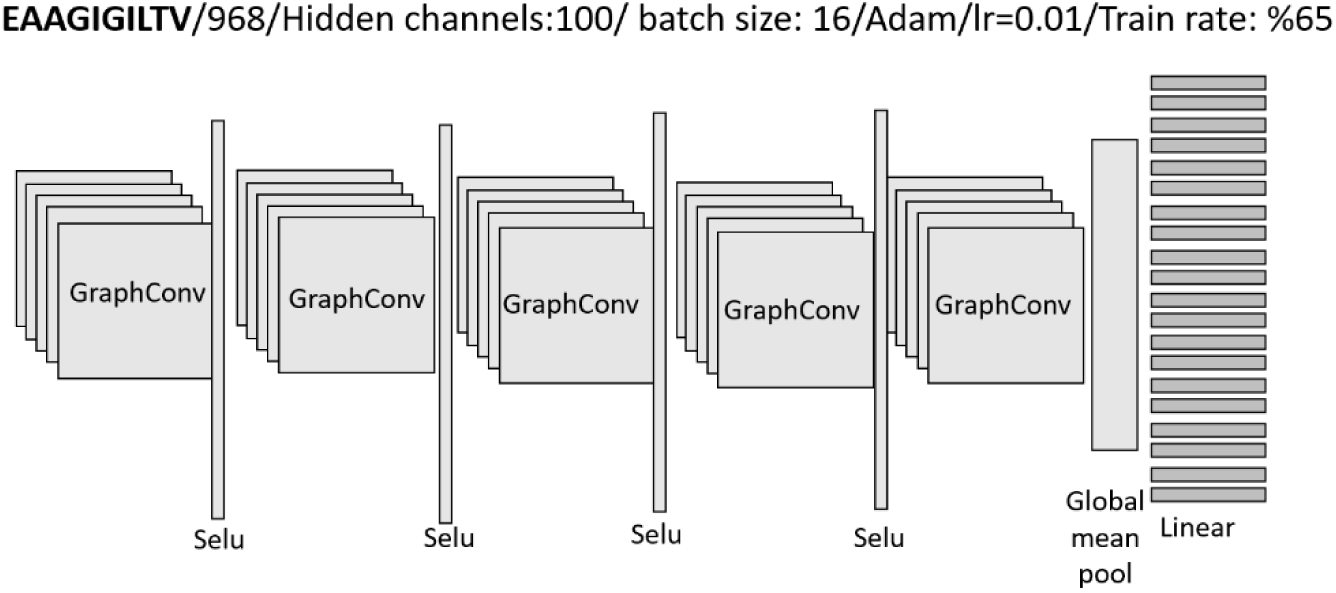
Model used for EAAGIGITV-TCRs binding

**Figure 8:**
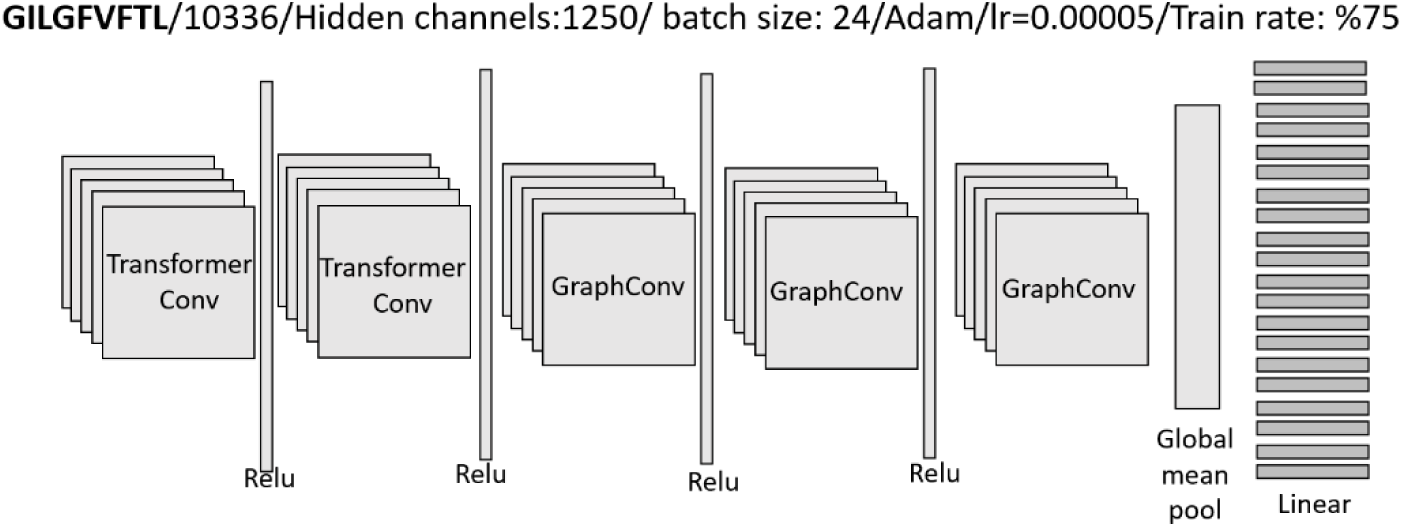
Model used for GILGFVFT-TCRs binding

**Figure 9:**
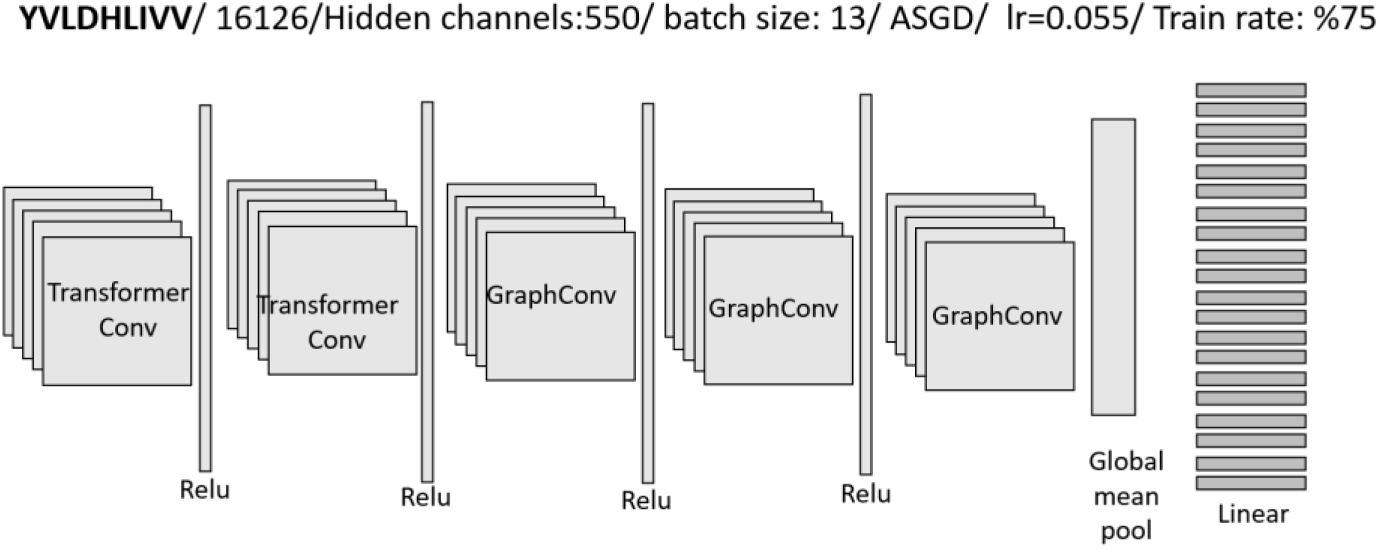
Model used for YVLDHIVV - TCRs binding

**Figure 10:**
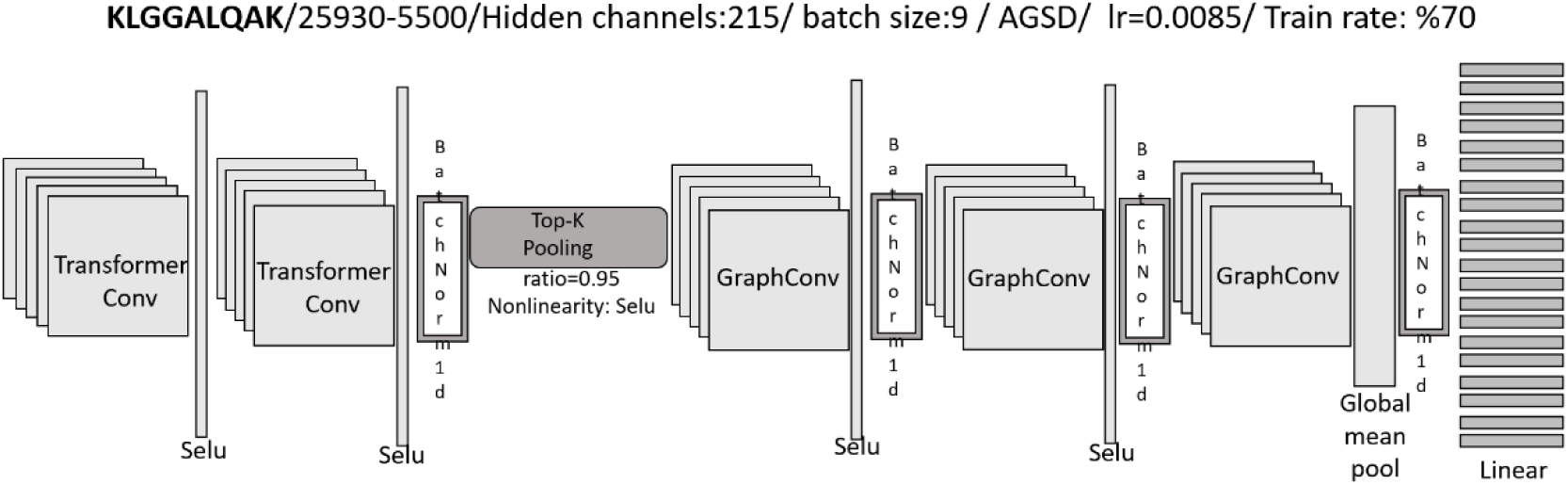
Model used for KLGGALQAK - TCRs binding

**Figure 11:**
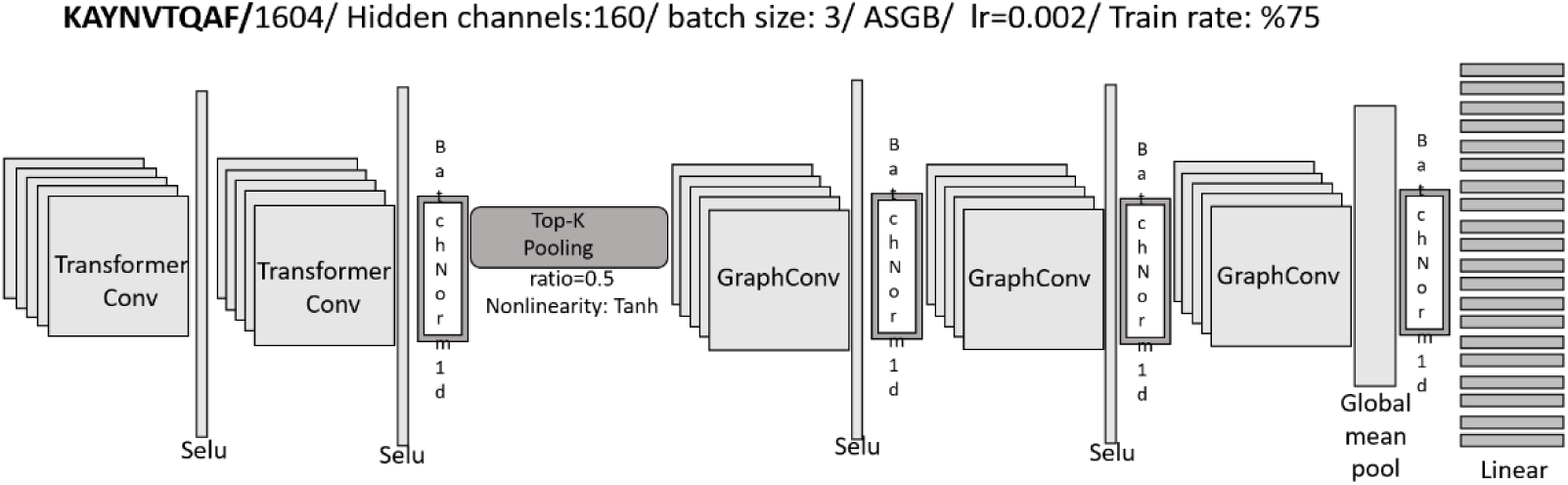
Model used for KAYNTVQAF - TCRs binding

The preceding models display variability in feature count, batch size, and neural network architecture, underscoring the distinct structural properties of each pairing.

### 5.3 Implementation and hyper-parameter tuning

For the epitopes KLGGALQAK and KAYNTVQAF, the datasets were substantially larger than those for other epitopes. However, when half of the data was assigned a label of 0 and the other half a label of 1, the models’ learning efficacy was compromised. A 50% ROC score indicated that the models indiscriminately assigned either the 0 or 1 label to all data points. The high degree of similarity between the amino acid chains labeled 0 and those labeled 1 revealed the models’ inability to differentiate between the two. This scenario represents a common challenge encountered in laboratory settings, where TCRs with similar sequences may either bind or not bind. To mitigate this issue, a new methodology was developed, employing ensemble learning. This technique, commonly applied in economics, exhibits the capacity to learn from datasets primarily comprising labels of 1 (Dong et al., 2020). In this study, the SMOTE method was implemented to generate synthetic labels of 0 (representing nonbinding) for the data labeled 1 (representing binding) (Chawla et al., 2002). As illustrated in Figure 12, for the complex red-blue dataset, synthetic data labeled 0 (shown in green) was generated for the data labeled 1 (shown in red) using the KNN algorithm. The resulting red-green dataset achieved a 98% ROC score with a machine learning model, demonstrating a high degree of discriminative power between the real data labeled 1 and the synthetic data labeled 0.

**Figure 12:**
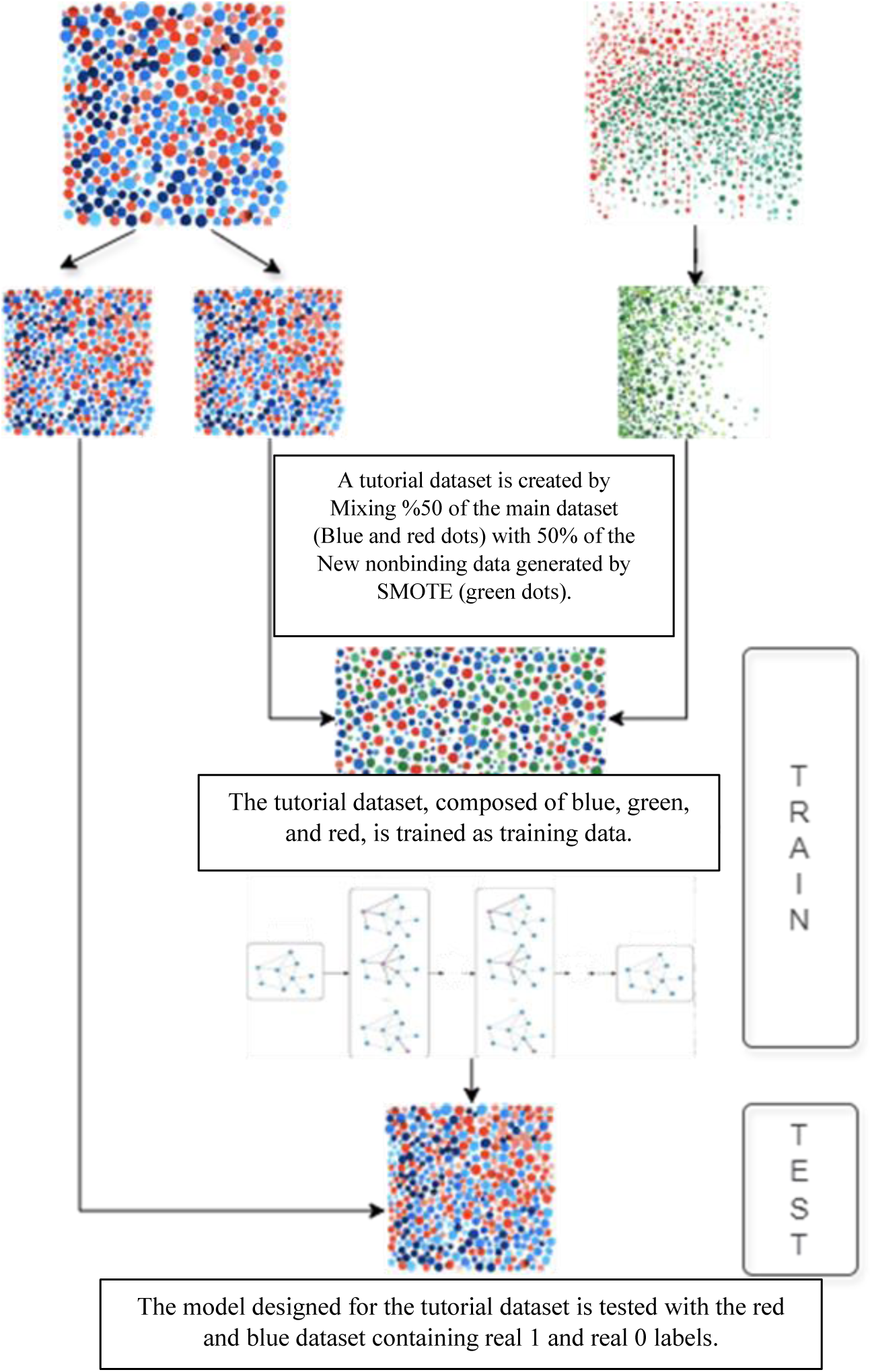
Training process of the model with the tutorial dataset

For model training, a subset of the red-blue dataset, comprising both real positive and negative labels, was used. To enhance training efficacy, a comparable quantity of synthetic negative labels was incorporated into the existing real negative labels. This augmentation created a hybrid dataset intended to improve the similarity between a portion of the real negative labels and the synthetic negative labels. This methodology enabled the model, which previously exhibited limitations in distinguishing between samples within the red-blue mixture, to achieve effective differentiation within the augmented red-blue-green dataset.

The trained model underwent testing utilizing the red-blue dataset, serving as the primary dataset. This dataset comprises empirically derived 0 and 1 labels, initially segregated, with subsequent reporting of model performance. This Tutorial Learning methodology yielded Receiver Operating Characteristic (ROC) scores of 71% and 67% for the KLGGALQAK and KAYNTVQAF peptides, respectively. The equilibrium between empirical 0 and synthetic 0 labels within the training dataset demonstrably influenced test performance metrics. Consequently, a strict 50-50 ratio is not a prerequisite for future investigations. A further critical parameter is the k value employed in the k-Nearest Neighbors (KNN) algorithm for synthetic 0 label generation. Empirical observations suggest that increased k values correlate with enhanced test performance, particularly for more complex datasets.

All experiments were conducted using a fixed random seed for reproducibility. To assess variance, we performed 5 independent runs with different random seeds and report the mean and standard deviation for each metric.

### 5.4 Evaluation metrics

All datasets comprised examples labeled 0 and 1, with each dataset containing an equal number of examples for each label, thereby establishing a balanced dataset for predictive modeling. Model performance was evaluated using metrics such as Accuracy, Precision, Recall, F1-score, and Area Under the Curve (AUC). These metrics are detailed below.

Accuracy serves as a crucial metric for evaluating the performance of a classification model. It quantifies the proportion of correctly classified instances relative to the total number of instances. This metric is particularly informative for summarizing overall model performance on datasets exhibiting class balance. True Positives (TP) denote instances correctly classified as positive, while True Negatives (TN) indicate instances correctly classified as negative.

Conversely, False Positives (FP) represent instances incorrectly classified as positive, and False Negatives (FN) denote instances incorrectly classified as negative. The calculation of accuracy is presented as

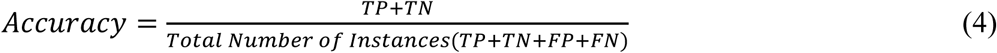

Precision represents the proportion of correctly predicted positive observations among all predicted positive observations, as expressed in Equation 5. Recall denotes the proportion of correctly predicted actual positive observations, calculated as shown in Equation 6. The F1 Score provides a balanced measure of precision and recall, defined as the harmonic mean of these two metrics, as demonstrated in the subsequent equations.

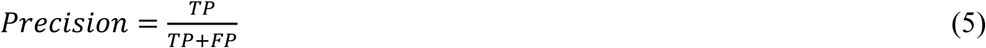

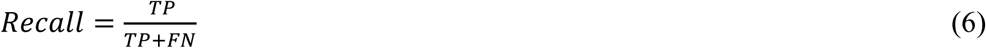

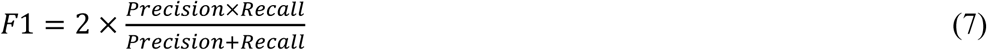

The Area Under the ROC Curve (AUC) is a frequently employed metric for evaluating the performance of binary classification models. This metric represents the area under the Receiver Operating Characteristic (ROC) curve, which illustrates the relationship between the True Positive Rate (TPR) and the False Positive Rate (FPR) across a range of threshold values. The AUC score provides a measure of the model’s ability to discriminate between positive and negative classes. The formulas for TPR and FPR are presented below:

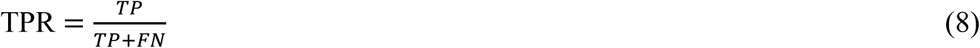

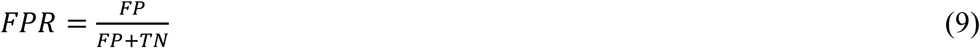

## Acknowledgment

Ethical and publication information related to the study is shared below.

<u>Epitope Data Source:</u>

- Smirnov AS, Rudik AV, Filimonov DA, Lagunin AA. TCR-Pred: A new web-application for prediction of epitope and MHC specificity for CDR3 TCR sequences using molecular fragment descriptors. Immunology. 2023; 169(4): 447–53. https://doi.org/10.1111/imm.13641. Source of the EAAGIGILTV dataset used in the above study: https://www.way2drug.com/TCRpred/definition.php
- Cai, M., Bang, S., Zhang, P., & Lee, H. (2022). ATM-TCR: TCR-epitope binding affinity prediction using a multi-head self-attention model. Frontiers in immunology, 13, 893247. https://doi.org/10.3389/fimmu.2022.893247. Source of the EAAGIGILTV, GILGFVFTL, YVLDHLIVV and KLGGALQAK dataset used in the above study: https://github.com/Lee-CBG/ATM-TCR
- Chen, J., Zhao, B., Lin, S., Sun, H., Mao, X., Wang, M., … & Xiong, Y. (2024). TEPCAM: Prediction of T-cell receptor–epitope binding specificity via interpretable deep learning. Protein Science, 33(1), e4841. https://doi.org/10.1002/pro.4841. Source of the KAYNTVQA dataset used in the above study: https://github.com/Chenjw99/TEPCAM

## Author information

### Author contribution

[1st Author Kenan Menguc]: Contributed significantly to data collection and analysis, took a leading role in the design and writing of the manuscript. Also contributed to the literature review and analysis, and made important edits during the writing process of the article.

[2nd Author Alper Yilmaz]: Played an important role in the idea development phase of the manuscript and made significant contributions to the writing process. Actively participated in the directed criticism and revision of the manuscript, thereby improving the quality of the final version. Additionally, contributed significantly to the critical revision and editing of the manuscript.

### Data availability

The dataset used in the study is open source and the study complies with ethical rules. The coding and dataset can be accessed from “ https://github.com/kenanmenguc/TCR-Epitope--Tutorial-Model “ as open source.

### Declarations

Competing interests the authors declare that they have no competing interests. Ethical approval and consent to participate not applicable. Consent for publication all data contained in the paper have the consent of all authors to be published. This research don’t involving human participants and/or animals.

## Footnotes

1 https://github.com/kenanmenguc/TCR-Epitope--Tutorial-Model

